# Fish primary embryonic stem cells self-assemble into retinal tissue mirroring *in vivo* early eye development

**DOI:** 10.1101/2021.01.28.428593

**Authors:** Lucie Zilova, Venera Weinhardt, Tinatini Tavhelidse, Thomas Thumberger, Joachim Wittbrodt

## Abstract

Organoids derived from pluripotent stem cells promise the solution to current challenges in basic and biomedical research. Further progress and widespread applications are however limited by long developmental time, variable success, and lack of direct comparison to an *in vivo* reference. To overcome those limitations, we derived organoids from rapidly developing teleosts. We demonstrate how primary embryonic stem cells from zebrafish and medaka efficiently self-organize into anterior neural structures, particularly retina. Within four days, blastula-stage cell aggregates reproducibly execute key steps of eye development: retinal specification, morphogenesis and differentiation. The number of aggregated cells as well as genetic factors crucially impacted upon the concomitant morphological changes that were intriguingly reflecting the *in vivo* situation. High reproducibility and rapid development of fish-derived organoids in combination with advanced genome editing techniques immediately allow addressing aspects of development and disease, and systematically probing the impact of the physical environment on morphogenesis and differentiation.

## Introduction

Mouse and human embryonic stem cells have been shown to self-assemble into retinal tissue when aggregated and cultured under 3D suspension culture conditions (Eiraku et al., 2011; Kuwahara et al., 2015; Nakano et al., 2012). Embryonic stem (ES) cells derived from teleosts (medaka and zebrafish) blastula-stage embryos contribute to all germ layers of chimeric embryos underlining their pluripotency (Ho et al., 2014; Hong et al., 1998, 1996; Peng et al., 2019; Robles et al., 2011; Yi et al., 2010). It has recently been shown that zebrafish blastula explants can form polarized structures that recapitulate some aspects of embryonic patterning and morphogenesis when cultured in the absence of yolk (Fulton et al., 2020; Schauer et al., 2020). However, under no condition reported so far, fish-derived explants or re-aggregates have formed highly organized neural structures.

## Results

### Generation of fish stem cell-derived aggregates

Here, we used primary embryonic stem cells derived from blastula-stage embryos of medaka (*Oryzias latipes*) as a source of pluripotent cells and established the conditions to generate the anterior neural structures, particularly retinal tissue. Previous studies performed with mouse and human ES cell aggregates have shown that low serum concentration in combination with 3D suspension culture support retinal specification and differentiation. In particular, addition of extracellular matrix components such as laminin-rich Matrigel promotes retinal formation (Eiraku et al., 2011; Nakano et al., 2012). We dissociated blastula-stage embryos and used U-shaped low adhesive wells to re-aggregate 1,000–2,000 cells in basic media supplemented with 5% KSR (knockout serum replacement) (Figure 1a, b). After one day of culture, media were supplemented with 2% Matrigel (Figure 1a). This protocol facilitates highly efficient aggregation and compaction of ES cells and resulted in a smooth and compacted morphology by day 1 (Figure 1b, c; Video 1). The aggregates formed an organized and layered structure that was apparent in the periphery of the aggregate from day two onwards (Figure 1c, d). Based on the higher order epithelial organization, we call the structures from day 2 on - organoids. The potential of blastomeres to form organoids was restricted to a blastula stage (1,000 – 2,000 cells) and cells derived from earlier developmental stages such as early and late morula failed to form stable aggregates or aggregated only partially (Figure S1). Thus, the exact time point of cell dissociation plays a decisive role for the success and efficiency of aggregation.

**Figure 1.**
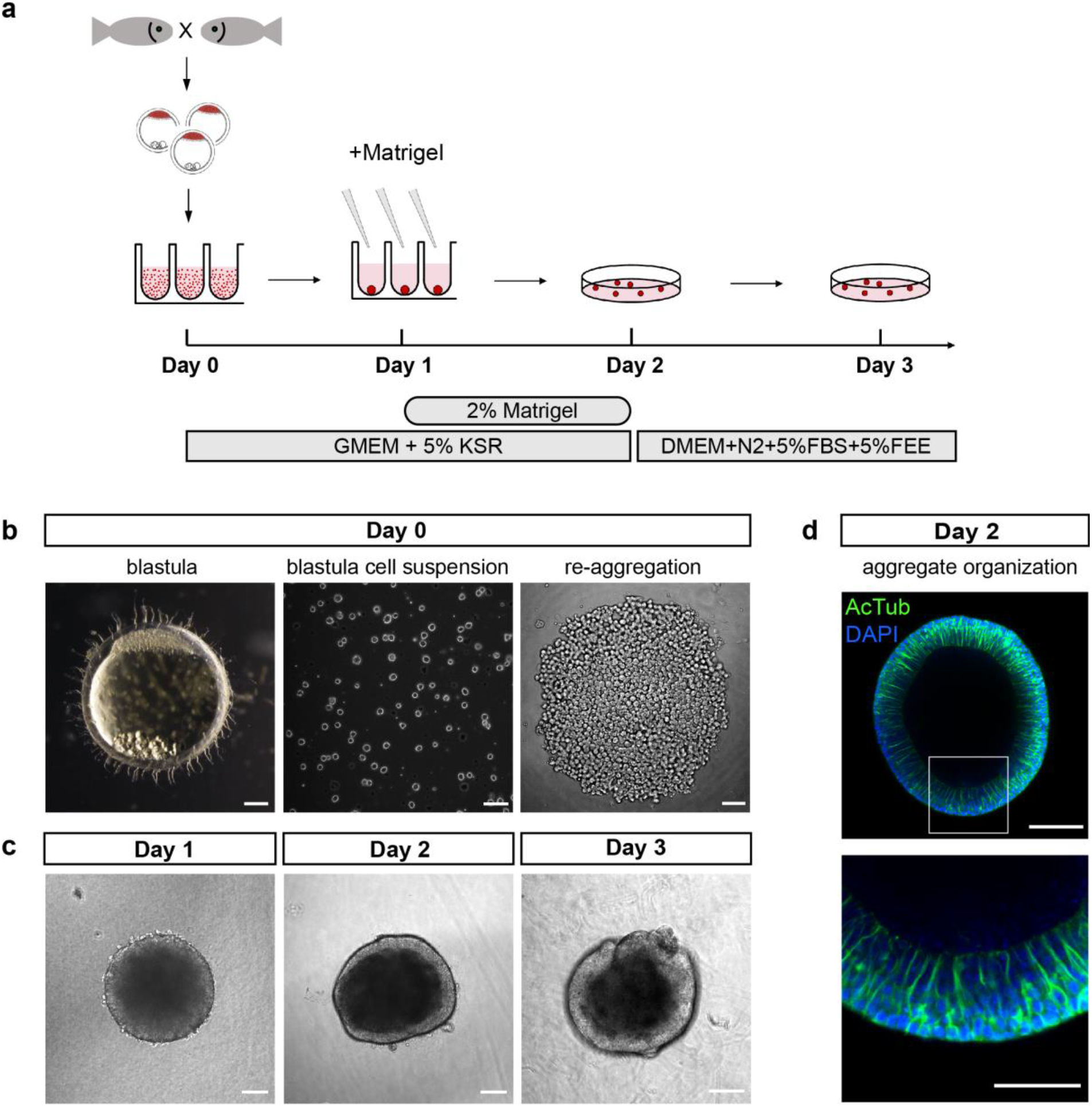
Generation of fish stem cell-derived aggregates. **(a)** Schematic representation of aggregate generation, its time line and culture conditions. At day 0, primary stem cells were harvested from blastula-stage medaka embryos and re-aggregated in low binding U-shape 96 well plates. At day 1, culture media was supplemented with Matrigel. At day 2, aggregates were transferred to a low binding culture plate and maintained in 3D suspension culture conditions in DMEM/F12 media supplemented with 5% FBS, 5% FEE and N2 supplement. Gross morphology of aggregates was analyzed at days 1, 2 and 3. KSR, knockout serum replacement; FBS, fetal bovine serum; FEE, fish embryonic extract. **(b)** Dark-field images of blastula-stage embryo, blastula-derived primary stem cell suspension and re-aggregated cells. **(c)** Gross morphology of aggregates at day 1, day 2 and day 3 after re-aggregation. **(d)** Optical section of aggregate organization visualized by staining against acetylated tubulin (AcTub) and DAPI nuclear stain Scale bars: 100 μm and 50 μm (enlargement in (d)).

### Fish-derived organoids form retinal neuroepithelium under the control of *Rx3*

Retinal specification in vertebrates is governed by the action of retina specific transcription factors such as Rx, Pax6, Six3, Sox2, Six6 and Lhx2 within the anterior neural plate (Li et al., 1997; Loosli et al., 1999; Zuber, 2003) and the formation of the optic vesicle neuroepithelium (Chow and Lang, 2001; Fuhrmann, 2010; Martinez-Morales et al., 2017). *Rx* genes, in teleosts represented by three paralogous genes (*Rx1, Rx2* and *Rx3*), are the earliest genes expressed by the retinal lineage (Chuang et al., 1999; Deschet et al., 1999; Loosli et al., 2003, 2001). To monitor the efficiency of retinal fate acquisition, we took advantage of the earliest expressed *Rx* paralogue, *Rx3* and employed a transgenic reporter line (*Rx3::H2B-GFP*) in which nuclear GFP is expressed under the control of the *Rx3* regulatory elements (Figure 2a, b) (Rembold et al., 2006). *Rx3::H2B-GFP* drives the expression of H2B-GFP already in the anterior neural plate and is subsequently found in the optic vesicle neuroepithelium and prospective forebrain of fish embryos at the first day post-fertilization (1 dpf) (Figure 2a). We addressed the onset of GFP expression in *Rx3::H2B-GFP* aggregates by time-lapse imaging over 18 h post-aggregation (hpa) and compared it to corresponding *Rx3::H2B-GFP* embryos for reference (Figure 2c, d; Video 2). All aggregates started expressing GFP at 15.5 ± 0.36 hpa (n = 17) with GFP expression in control *Rx3::H2B-GFP* embryos starting at 13.5 ± 1.03 h (n = 6) after blastula stage (Figure 2d), corresponding to the onset of *Rx3* expression at around 20 hpf (Loosli et al., 2001). In comparison to the embryo, the organoids showed a delay of GFP expression of approximately three hours which can be attributed to the time of re-aggregation (Video 1). *Rx3* expression was initiated already at day 0 in the absence of Matrigel, indicating that the acquisition of retinal fate occurs spontaneously and independently of extracellular matrix components.

**Figure 2.**
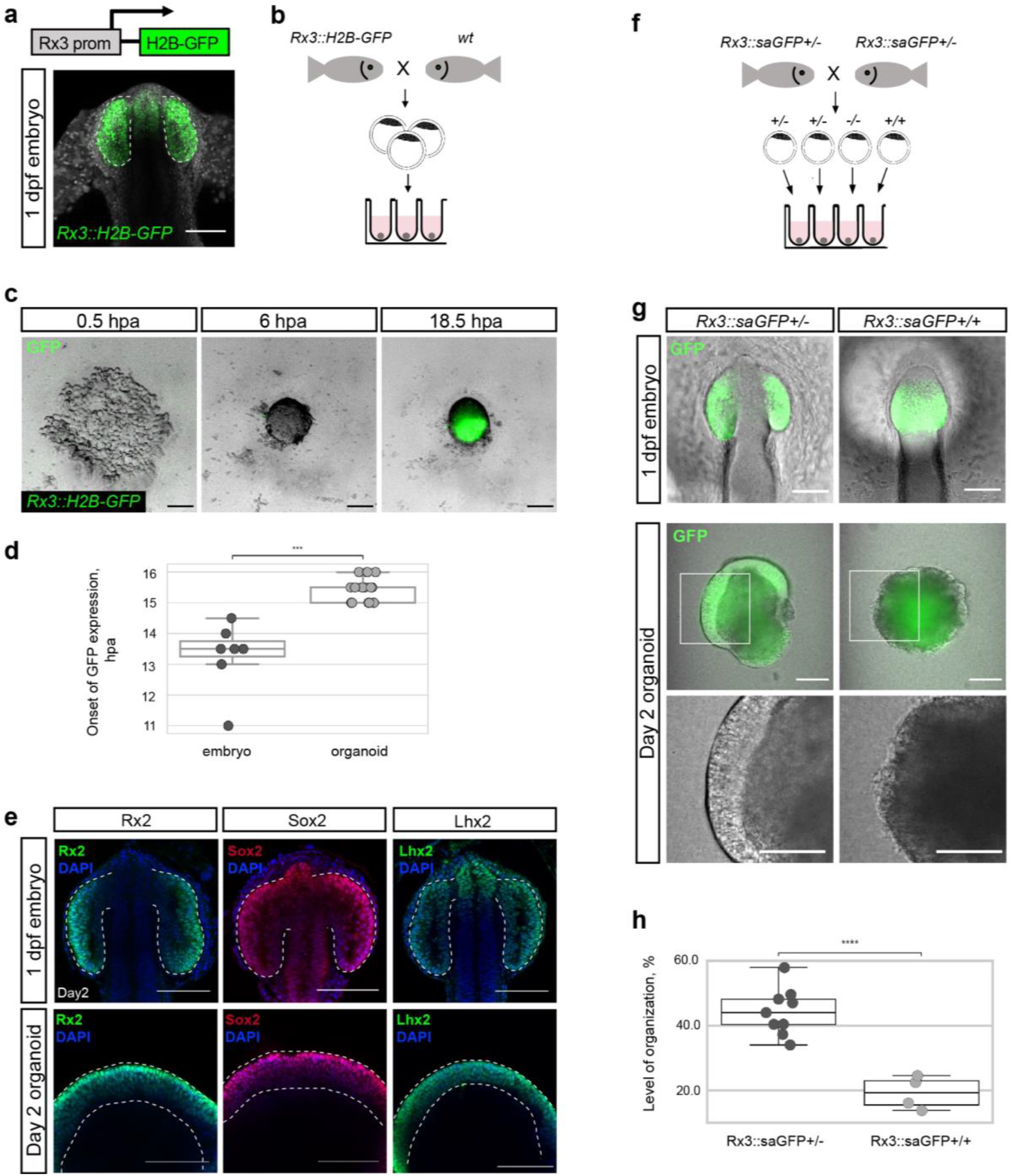
Fish-derived organoids form retinal neuroepithelium under the control of *Rx3*. **(a)** Schematic representation of *Rx3::H2B-GFP* transgenic construct and corresponding expression domain of GFP in optic vesicles of developing medaka embryo at 1 dpf. **(b)** Scheme of organoid generation from *Rx3::H2B-GFP* transgenic fish. **(c)** Bright-field and fluorescence images of aggregates derived from *Rx3::H2B-GFP* transgenic fish at 0.5, 6 and 18.5 hpa. **(d)** Analysis of the onset of GFP expression in *Rx3::H2B-GFP*-derived fish embryos (n = 6) and organoids (n = 17). ^∗∗∗^p<0.001. **(e)** Optical sections of embryonic heads (upper panel) and organoids (lower panel) stained with antibodies showing expression of Rx2, Sox2 and Lhx2 co-stained with DAPI nuclear stain. Dashed lines indicate optic vesicle in embryo or neuroepithelium in organoid. **(f)** Scheme of aggregate generation formation from *Rx3*-deficient (*Rx3::saGFP*) single blastulae. **(g)** Bright-field and fluorescence images of phenotypes of Rx3 +/-heterozygote and *Rx3*-/-mutants at 1dpf and corresponding organoids at day 2. **(h)** Analysis of the organization level in organoids, measured as % of area occupied by neural epithelium to area of organoid, in *Rx3*-deficient (*Rx3::saGFP*) aggregates (n = 13). ^∗∗∗∗^p<0.0001. dpf, days post-fertilization; hpa, hours post-aggregation; hpf, hours post-fertilization; wt, wild-type. Scale bars: 100 μm.

To address whether GFP-positive aggregates followed retinal specification, we analyzed the expression of retinal markers expressed at subsequent stages of retinal development. The expression of the transcription factors Rx2, Sox2 and Lhx2 (Figure 2e) in the epithelium of day 2 organoids resembled their expression in the optic vesicle neuroepithelium at 1 dpf and underlines that the aggregates adopted retinal fate. All organoids analyzed (n ∼1,500) from 52 independent experiments formed retinal neuroepithelia indicating that retinal specification under the established conditions is very robust and highly efficient.

The formation of the optic vesicle neuroepithelium strictly depends on expression of the transcription factor Rx such that in *Rx3*-null mutants, formation of the optic vesicle is disrupted leading to the eyeless phenotype (Loosli et al., 2003, 2001; Mathers et al., 1997; Rembold et al., 2006; Stigloher et al., 2006). Organoids derived from *Rx3*-null mutants could be used to investigate the failure of retinal neuroepithelium formation. Unfortunately, null alleles resulting from mutagenesis screens do not contain traceable markers. We therefore envisioned to perform a targeted gene-trapping approach by inserting an open reading frame-adjusted gene trap cassette comprising a splice acceptor and a *GFP* sequence (saGFP) for visual readout followed by a polyA and strong terminator sequence derived from the ocean pout (OPT; (Clark et al., 2011, Figure S2a)). Using the CRISPR/Cas9 system for such targeted integration, we generated traceable *Rx3* mutants (*Rx3::saGFP*; Figure S2a). While *Rx3::saGFP* heterozygotes develop normally and give rise to GFP labelled optic vesicles, homozygous mutant embryos fail to evaginate the optic vesicle from the lateral wall of the diencephalon (Figure 2g, S2b). Interestingly and in contrast to the *eyeless* mutant described (Loosli et al., 2001), this inability to organize the retinal neuroepithelium was not temperature sensitive. To address whether fish-derived organoids retain genetic constraints and the developmental program displayed in the embryo, we derived organoids from individual *Rx3*-/- and Rx3 +/-blastulae (Figure 2f). Like in the embryonic context, the retinal neuroepithelium was present in the Rx3+/-(n = 9) organoids by day 2. Aggregates derived from individual *Rx3*-mutant homozygote blastulae (*Rx3*-/-) (n = 4) showed low level of organization, failing to form retinal neuroepithelium by day 2 (Figure 2g, h). This data indicate that the formation of retinal neuroepithelium is hard-wired also in the organoid and follows the same developmental program as *in vivo*, also upon loss-of-function. (Loosli et al., 2001; Rembold et al., 2006; Stigloher et al., 2006; Winkler et al., 2000).

We next asked whether spontaneous acquisition of retinal fate is generally applicable to other fish species, for example the evolutionarily distant zebrafish (*Danio rerio*). We used zebrafish blastulae as a source of pluripotent stem cells to generate retinal organoids. To monitor the acquisition of retinal fate, we used the expression of a nuclear GFP controlled by the regulatory element of the early retinal marker gene *Rx3*. We used a transient assay and injected the *Rx3::H2B-GFP* reporter DNA into one cell stage embryos and generated cell aggregates when embryos reached blastula stage (Figure 3a). Zebrafish cells aggregated as efficiently as cells from medaka and showed epithelial organization already by day 1 (Figure 3b). Considering the faster development of zebrafish compared to medaka (Iwamatsu, 2004; Kimmel et al., 1995), we asked whether the difference in organismal development is reflected in self-assembly and organization of aggregates. As for medaka, we performed time-lapse imaging of GFP expression in zebrafish aggregates over 18 hpa and compared it to the corresponding *Rx3::H2B-GFP* reporter fish as reference (Figure 3c; Video 3). Zebrafish embryos started to express GFP at 7.88 ± 0.77 h (n = 6) after blastula stage (corresponding to ∼10 hpf), 8 h earlier than medaka *Rx3::H2B-GFP* embryos. Aggregates generated from zebrafish embryos formed organoids that started expressing GFP at 9.75 ± 0.21 hpa (n = 15), following their endogenous timer with a delay of 3 h in *Rx3* expression as observed for medaka. Similar to medaka we attribute this delay to the time the cells take to re-aggregate. Thus, the formation of the neuroepithelium in aggregates reflects the species-specific difference in the relative pace of development.

**Figure 3.**
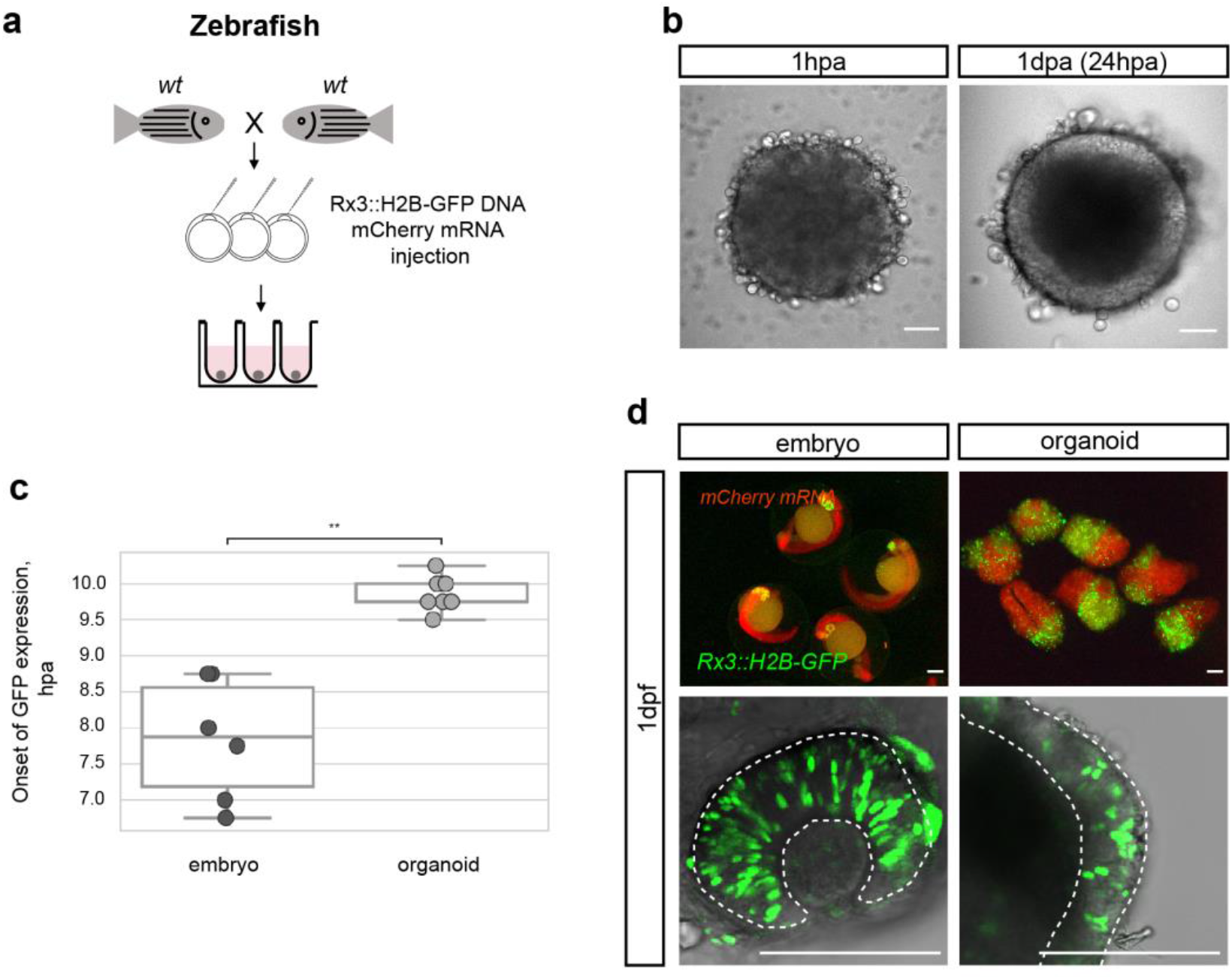
Zebrafish blastula-derived primary embryonic stem cells self-organize into retinal organoids. **(a)** Scheme of aggregate generation from *Rx3::H2B-GFP* DNA-injected blastulae. **(b)** Bright-field images of zebrafish-derived aggregates at 1 hpa and 1 dpa. **(c)** Analysis of the onset of GFP expression in *Rx3::H2B-GFP*-derived zebrafish embryos (n = 6) and organoids (n = 15). ^∗∗^p<0.01 **(d)** Fluorescent images of mCherry and GFP expression domains in *Rx3::H2B-GFP* injected embryos at 1 dpf and corresponding aggregates at day 1. Dashed lines indicate the embryonic retina and organoid retinal neuroepithelium, respectively. wt, wild-type; dpf, days post-fertilization; dpa, days post-aggregation; hpa, hours post-aggregation. Scale bars: 100 μm.

*Rx3* reporters for retina formation (*Rx3::H2B-GFP*) displayed H2B-GFP expression in the developing retina at 1 dpf and aggregates formed organoids showing retinal identity as indicated by *Rx3* expression in the outer cellular layer (Figure 3d) by day 1. This data show under the established conditions fish primary embryonic stem cells to differentiate into retinal neuroepithelium, a feature found over a wide evolutionary distance from medaka to zebrafish suggestive of deep conservation of early retinal development, irrespective of the embryonic environment.

### Fish organoids form optic vesicle-like structures

Development of a functional retina is accompanied by several morphological events that ultimately result in the formation of a functional retina. One of them is the formation of the optic vesicle. In most fish-derived organoids generated by aggregation of 1,000–2,000 cells (corresponding to the approximate size/number of cells of a single blastula), optic vesicle formation was not favored (Figure 4a, b). These “regular size” organoids formed a continuous neuroepithelium on their entire surface (Figure 1c, d). It has been previously reported that in some *in vitro* self-organizing systems the number of interacting cells can generate a bias towards particular morphological processes (Brink et al., 2014; Fulton et al., 2020; Völkner et al., 2016). Thus, we reduced cell-seeding density by 50% and analyzed the expression patterns of retinal-specific markers (Rx2 and *Rx3::H2B-GFP*). Interestingly, small organoids (generated by aggregation of 500–800 cells) showed expression of Rx2 and *Rx3* only in restricted regions (Figure 4a, b). Regular size aggregates (1,000–2,000 cells) produced single uniform domains of retinal neuroepithelium in all experiments, while small aggregates tended to form one to four isolated optic vesicle-like retinal domains (54% one domain, 40% two domains, 5% three domains and 1% four domains) (Figure 4b–c, S3). The size of these isolated retinal domains in small aggregates (154.5 ± 28.6 μm; n = 56) was reminiscent of the optic vesicle in embryos at 1 dpf (162 ± 10.6 μm; n = 16) (Figure 4d).

**Figure 4.**
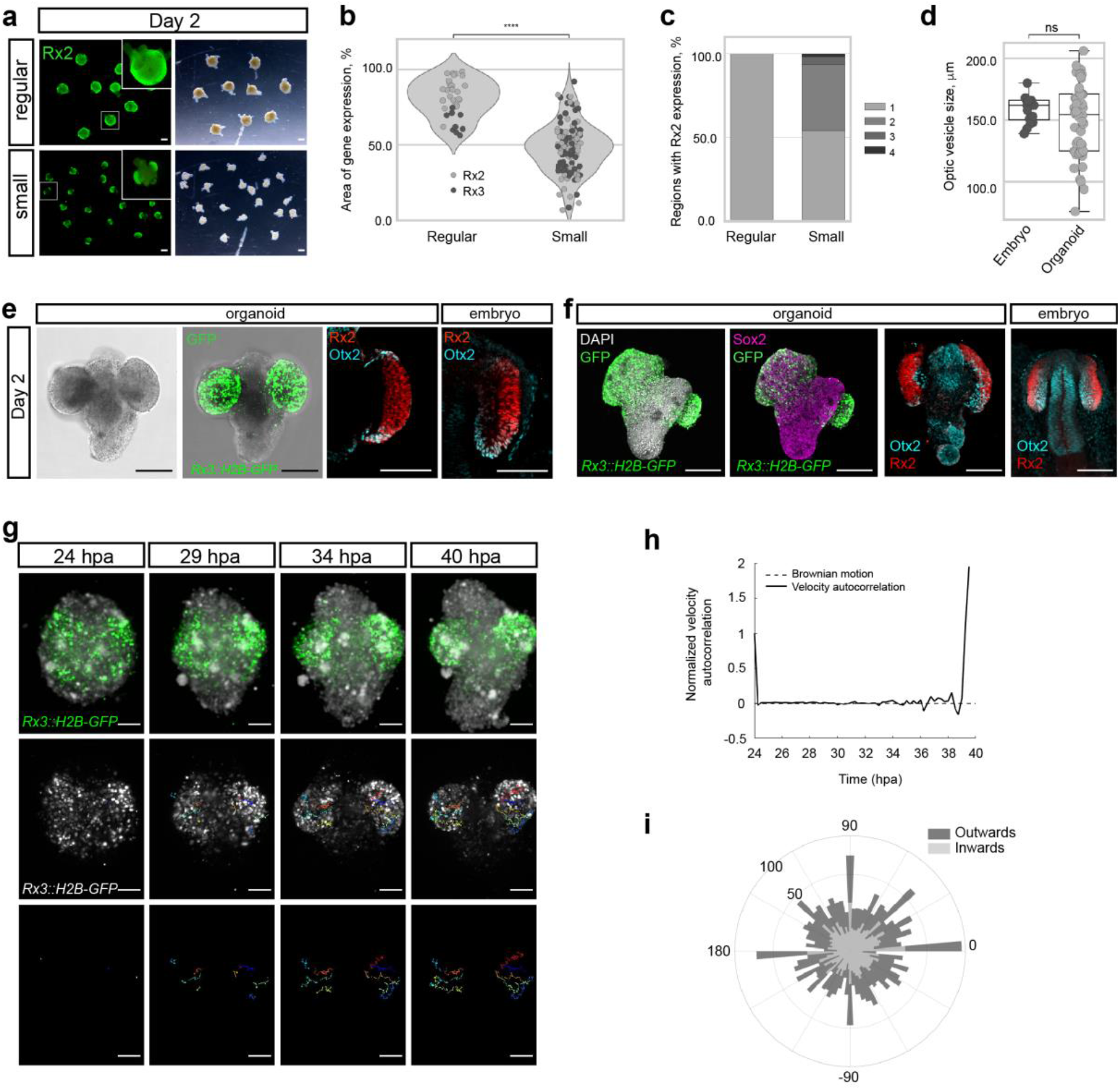
Self-organization of primary embryonic stem cells into optic vesicle-like structures. **(a)** Fluorescence and bright-field images of day 2 organoids produced with regular (1,000–2,000) and small (500–800) cell-seeding densities stained with anti-Rx2 antibody. **(b)** Analysis of the area of Rx2/*Rx3* expression area (% of total organoid area) in day 2 organoids. ^∗∗∗∗^p<0.0001. **(c)** Distribution of individual gene-expressing domains (1–4) per organoid produced with regular (n = 26 for Rx2, n = 9 for *Rx3*) and small (n = 57 for Rx2, n = 66 for *Rx3*) seeding densities from 9 independent experiments. **(d)** Size, measured as largest circumference, of the optic vesicle of 1dpf embryos and optic vesicle-like structures formed by day 2 organoids (n = 16 embryos, n = 56 aggregates from 6 independent experiments). ns, non-significant. **(e)** Bright-field and fluorescent images of day 2 *Rx3::H2B-GFP* organoids stained with anti-GFP antibody. Optical sections of organoid (day 2) and embryo (1 dpf) stained with anti-Rx2 and anti-Otx2 antibodies. **(f)** Maximal projection of day 2 organoids and 1 dpf embryo generated from *Rx3::H2B-GFP* transgenic or wild type blastulae and stained with neural tissue-specific anti-Sox2 and anti-Otx2 antibodies, co-stained with anti-Rx2 and DAPI nuclear stain. **(g)** *In vivo* time-lapse images acquired with MuVi SPIM of optic vesicle-like structure evagination in *Rx3::H2B-GFP* retinal organoids. From top to bottom: maximal projection of quasi bright-field, GFP and tracks of exemplary cells. **(h)** Normalized velocity autocorrelation for all tracks. **(i)** Directional histogram of tracks (n = 4600, from two optic vesicles) separated for inward and outward movements. hpa, hours post-aggregation; dpf, days post-fertilization. Scale bars: 100 μm.

Soon after evagination, two major domains are being specified within the optic vesicle. The distal/ventral region, expressing retina-specific genes, further differentiates into the retina, while the dorsal/outer region, expressing transcription factor Otx2, differentiates into pigmented epithelium (Bovolenta et al., 1997; Fuhrmann, 2010; Hatakeyama et al., 2001; Hirashima et al., 2008). Interestingly retinal domains produced by small organoids displayed optic vesicle-like morphology with specifically localized expression of Rx2 and Otx2 transcription factors, marking prospective retinal and retinal pigmented epithelium territories respectively (Figure 4e) indicating further compartmentalization of the organoid-derived optic vesicle.

Besides the retinal tissue, small aggregates also produced an Rx2-negative non-retinal domain expressing the general neuronal marker Sox2 (Pevny and Placzek, 2005; Wegner and Stolt, 2005) and the forebrain marker Otx2 (Acampora et al., 1995; Simeone et al., 1993; Tian et al., 2002), strongly indicating that the neighboring tissue adopted anterior neuronal identity (Figure 4f). In some organoids, intriguingly, retinal and non-retinal domains showed a complex architecture, resembling a “head with two eyes” (Figure 4e–g).

To address the mode of optic vesicle formation in fish organoids, we employed live imaging of *Rx3::H2B-GFP*-derived organoids from day 1 to day 2 of development. Figure 4g shows development of the organoid forming two opposing optic vesicle-like structures. At the initial time point (24 hpa), GFP-positive cells were distributed throughout the whole volume of the organoid indicating that cells acquired retinal identity individually. Single cell tracking and analysis of cell displacement showed that initially cells behaved as freely diffusing particles (Figure 4h). This behavior changed at about 34 hpa, when GFP-positive cells showed directed migration – non-linear growth of mean squared displacement – with an estimated speed of 0.413–0.479 μm/min resulting in the morphogenesis of the optic vesicle-like structures by 40 hpa (Figure 4g, h; Video 4). A similar transition from linear to nonlinear diffusion, due to increasing density of nuclear packing, was recently demonstrated *in vivo* in the growing retina of 24 hpf zebrafish (Azizi et al., 2020). Directionality analysis of single cell tracks (Figure 4i) showed that 51% of the Rx3-expressing cells were moving from the center to the periphery (outwards) of the organoid and contributed to the formation of the optic vesicle (see dark blue, red, orange and yellow tracks in Figure 4g). About 49% of the initially laterally located cells moved in the opposite direction – towards the interior of the organoid (inwards) (light green track in Figure 4g). These outward- and inward-directed cell movements were highly reminiscent of the migratory behavior of Rx3-expressing cells during optic vesicle formation in medaka fish (Rembold et al., 2006). Notably, a similar behavior was observed in the organoids forming single and multiple optic vesicles (Video 5). Altogether these results show, that the cell behavior in the embryonic stem cell-derived organoids closely mimics *in vivo* development.

It is worth mentioning that the organization of retina-committed cells into the optic vesicle neuroepithelium is strictly dependent on extracellular matrix components as *Rx3::H2B-GFP*-derived organoids failed to form optic vesicle structures and eventually died in the absence of Matrigel (Video 6). Accordingly, laminin-1 (the main component of the Matrigel) is highly enriched specifically around the basal surface of the forming optic vesicle and has been found to play an essential role in cell polarization and maintenance of neuroepithelial cell morphology during optic vesicle evagination (Ivanovitch et al., 2013).

### Fish organoids show onset of retinal differentiation

Retinal specification is followed by the process of differentiation which leads to the generation of seven retinal cell types (retinal ganglion cells, amacrine cells, horizontal cells, bipolar cells, rod and cone photoreceptors and Müller glia cells) organized in three nuclear layers. One of the first hallmarks of retinal differentiation is the expression of the transcription factor Atoh7 (Brown et al., 1998; Del Bene et al., 2007; Kay et al., 2001). Atoh7-positive progenitors have been found to give rise to retinal ganglion cells, amacrine cells, horizontal and photoreceptor cells during fish retinal development (Poggi et al., 2005). We therefore monitored *Atoh7* expression in fish-derived retinal organoids, using an *Atoh7::EGFP* transgenic line (Del Bene et al., 2007) (Figure 5a–c). In fish embryos, EGFP expression was located specifically in the differentiating retina at 2 dpf (Figure 5c). Day 3 organoids generated from *Atoh7::GFP* blastulae showed specific morphology with EGFP-negative non-retinal and EGFP-expressing retinal domains (Figure 5b; Video 7), indicating that the retinal differentiation program was successfully initiated. Furthermore, non-retinal regions proceeded through the process of neuronal differentiation as indicated by expression of HuC/D (Figure 5c; Video 7), a marker of early post-mitotic neurons (Good, 1995; Kim et al., 1996; Park et al., 2000).

**Figure 5.**
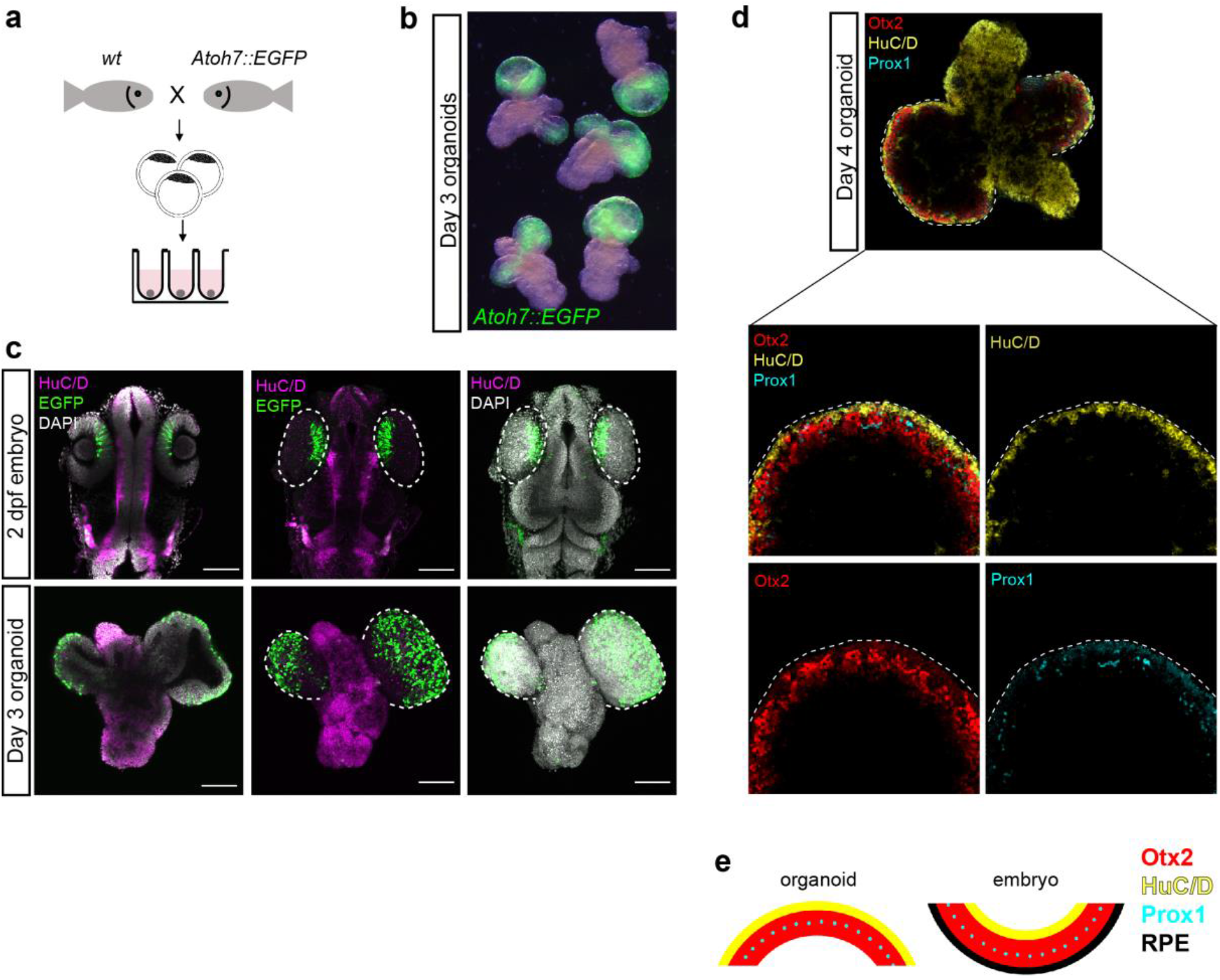
Onset of retinal differentiation. **(a)** Scheme of organoid generation from *Atoh7::EGFP* transgenic fish. **(b)** Fluorescent images of day 3 *Atoh7::EGFP* organoids. **(c)** Optical sections and maximal projections showing EGFP expression in the eye of the developing embryo at 2 dpf and retinal organoid at day 3 co-stained with antibody against HuC/D and DAPI. **(d)** Optical sections showing expression of HuC/D (amacrine and ganglion cells), Otx2 (bipolar and photoreceptor cells) and Prox1 (horizontal cells) in day 4 organoid. Dashed lines indicate the embryonic retina and organoid retinal domain. **(e)** Sketch of arrangement of cellular layers in organoid and embryonic retina. dpf, days post-fertilization. Scale bars: 100 μm.

Further analysis of the day 4 organoids showed that retinal domains consisted of differentiating retinal neurons organized in multiple cellular layers with cells differentiating towards amacrine and ganglion (expressing HuC/D) (Kay et al., 2001; Link et al., 2000; Park et al., 2000), photoreceptor and bipolar (expressing Otx2) (Fossat et al., 2007; Glubrecht et al., 2009; Koike et al., 2007; Nishida et al., 2003) as well as horizontal (expressing Prox1) (Dyer et al., 2003) cell lineages (Figure 5d). Interestingly, the arrangement of respective cellular layers was inverted when compared to normal retinal organization (Figure 5e). This can be attributed to the morphological differences, particularly optic vesicle to optic cup transition, between retina morphogenesis *in vivo* and organoid. In organoids, retinal differentiation is initiated without optic cup formation, most probably leading to the inverted retinal organization.

Taken together, our findings show that primary embryonic stem cells derived from blastula-stage fish embryos self-organize into aggregates. By low cell numbers and addition of Matrigel these aggregates are efficiently directed to retinal fate, through optic vesicle morphogenesis to retinal differentiation.

## Discussion

During embryogenesis, individual cells interact with each other and the environment to establish a 3D structures of high complexity, tissues, organs and ultimately the entire embryo, by a series of tightly controlled, highly stereotypical events. 3D *in vitro* cultures, organoids, from human and mouse embryonic stem cells have been established that provide a unique platform to study development as well the roots of pathologies under tightly controlled conditions (Brooks et al., 2019; Fligor et al., 2018; Gao et al., 2020; Kruczek and Swaroop, 2020; Kuwahara et al., 2015; Lancaster and Knoblich, 2014; Völkner et al., 2016). In the absence of immediate access to mammalian embryos, organoids derived from mammalian species offered to fill the gap. While human organoids are the most desired, they also take the longest time to develop and their systematic analysis is limited by technical challenges in genetic manipulations and, last not least, by the lack of direct comparison to the *in vivo* biological processes.

As all key steps of eye development are conserved across vertebrates, for decades the analysis in accessible genetic model systems had been the approach of choice. The derivation of organoids from fast developing non-mammalian models may similarly provide an easier and more immediate way to address different aspects of eye development.

Organoids from a wide range of vertebrate model organisms, such as medaka and zebrafish, hold the potential to elucidate evolutionary aspects of species diversity, key mechanisms underlying organogenesis and the etiology of disease. Key mechanisms can be directly compared to *in vivo* processes. Furthermore, the ease of highly efficient genome editing in fish (Gutierrez-Triana et al., 2018; Prykhozhij and Berman, 2018; Stemmer et al., 2015) in combination with rapid development, small size and full transparency offers an efficient technological platform for addressing diseases and testing therapies in large arrays, which then can be translated to mammalian species. Fish organoids may serve as “rapid prototyping system” to rapidly establish routes for the reliable control organogenesis even in the experimentally and temporally more demanding mammalian systems.

Although fish blastula-derived cells have been used to generate ES cultures (Ho et al., 2014; Hong et al., 1998, 1996; Peng et al., 2019; Robles et al., 2011; Yi et al., 2010), our current understanding of their ability to differentiate/assemble into particular tissues is very limited. Zebrafish blastula explants have been found to form polarized aggregates capable of specification of all three germ layers (Fulton et al., 2020; Schauer et al., 2020), resembling mammalian ES cell-derived gastruloids (Beccari et al., 2018; Brink et al., 2014; Moris et al., 2020; Turner et al., 2017). Besides that, fish primary embryonic stem cells have been directed to differentiate to cardiomyocytes forming beating cell sheets with contractile kinetics and electrophysiological features when cultured under defined chemical and mechanical conditions (Xiao et al., 2016). Here, we demonstrate that primary ES cells derived from medaka and zebrafish efficiently self-assemble into retinal organoids under culture conditions similar to those established for mammalian ES cells (Eiraku et al., 2011; Nakano et al., 2012). Our results show that certain developmental routes in culture are executed autonomously.

It had been previously proposed that anterior neural development represents the default differentiation program followed by pluripotent stem cells (Eiraku et al., 2011; Hemmati-Brivanlou and Melton, 1997; Kuwahara et al., 2015; Levine and Brivanlou, 2007; Nakano et al., 2012; Tropepe et al., 2001; Turner et al., 2014). In addition, inter-species blastula cell transplantation experiments between zebrafish and medaka result in ectopic retina formation composed exclusively of donor cells (Fuhrmann et al., 2020), further reflecting the intrinsic tendency of ES cells to form retinal structures. We found formation of fish retinal organoids highly efficient, reproducible and robust. We found that the starting size (cell number) of the initial aggregate crucially impacts on the morphogenesis of the resulting organoid. While small aggregates differentiated into anterior neuroectoderm that subsequently underwent morphogenesis bulging out retinal primordia that closely resembled optic vesicles, large aggregates entirely differentiated into a single giant retina. This appeared counterintuitive, since the “classical” reaction-diffusion model (Meinhardt, 2012; Turing, 1952) predicts that the number of repeating patterns – optic vesicles – will increase proportionally with tissue size. For fish retinal organoids, the effect is however inverted posing it as a model to address alternative pattering scenarios.

Re-aggregation of dissociated fish ES cells, was fast (about 2–3 h) and highly efficient when derived from blastula stage embryos. With optic vesicle formation by day 2 and the onset of differentiation by day 3, fish organoids proceed through development almost ten times faster than their mammalian counterparts (Eiraku et al., 2011; Kuwahara et al., 2015; Nakano et al., 2012). The formation of fish retinal organoids follows the *in vivo* developmental timing, reflecting the difference in relative speed of development of corresponding species.

Although eye development is highly stereotypic across vertebrates and the players involved are highly conserved, there are certain species-specific features that are also manifested in organoids. One such example is the cellular mechanism of optic vesicle evagination. In mammalian eye development, paralleled in mouse and human retinal organoid cultures, the optic vesicle is formed by a set of coordinated collective epithelial cell movements driving vesicle evagination from the initially formed neuroepithelium (Chauhan et al., 2015; Eiraku et al., 2011; Martinez-Morales et al., 2017). In teleosts, in contrast, optic vesicle formation is driven by the individual migration of Rx3-expressing retinal progenitor cells (Rembold et al., 2006). Fish-derived retinal organoids exhibit the same mechanism of optic vesicle formation, as recapitulated by the migratory trajectories and individual movements of *Rx3*-expressing cells (Figure 4g, j). Our analysis indicates that this fundamental property does not depend on the environment (embryo vs culture) and rather reflects a “hard wired” feature of self-organization. While the concept of evolutionary conservation allows to deduce fundamental mechanism by comparison of embryonic stages and the crucially contributing corresponding genes, this is not immediately transferable to the comparison of organoids from different species. Selection acts only on development and consequently relies on the entire context of the developing embryo. Morphological and molecular resemblances of organoids to the organismal counterparts indicate on the one hand their self-organizing capacity, irrespective of the environment or, alternatively may reflect that the conditions chosen mimic the organismal constraints. The comparison of organoids derived from a wide range of evolutionarily diverse species provides the opportunity to systematically address and distinguish intrinsic and extrinsic constraints contributing to organ morphogenesis and self-organization to ultimately tackle the question of the “conservation” of cellular self-organization, a feature that has apparently not been selected for in evolution.

## Materials and Methods

### Fish handling and maintenance

Medaka (*Oryzias latipes*) and Zebrafish (*Danio rerio*) stocks were maintained according to the local animal welfare standards (Tierschutzgesetz δ11, Abs. 1, Nr. 1, husbandry permit AZ35-9185.64/BH, line generation permit number 35–9185.81/G-145/15 Wittbrodt). The following medaka lines were used in this study: Cab strain as a wild-type, *Rx3::H2B-GFP* (Rembold et al., 2006), *Atoh7::EGFP* (Del Bene et al., 2007), *Rx3::saGFP* (this study). The following zebrafish lines were used in this study: AB zebrafish as a wild-type.

### Cloning of pGGD(saGFP-OPT-MCS) and generation of Rx3::saGFP-OPT knock-out line

The ocean pout polyA terminator (OPT) sequence was released from plasmid GBT-RP2 (Clark et al., 2011) via restriction digest with BfaI-FD (Thermofisher Scientific), ligated into pGGEV_5_linker (Addgene #49285), and fused into pGGDestSC-ATG (#49322 Addgene) using the Golden GATEway cloning system (Kirchmaier et al., 2013) and the following sequences: target site sequence of sgRNA *GFP_T1* (Stemmer et al., 2015) inserted in pGGEV_1, a multiple cloning site in pGGEV_2, a strong AD splice acceptor (Centanin et al., 2011) fused with a *GFP* variant not targeted by sgRNA *GFP_T1* (Stemmer et al., 2015) in pGGEV_3, pGGEV_4_linker (#49284 Addgene), a multiple cloning cassette in pGGEV_6 and pGGEV_7’ _linker (#49293 Addgene) into. Adjustment of open reading frame following the splice acceptor was accomplished via Q5 (NEB) mutagenesis by inserting a single or two nucleotides (+1 or +2, respectively) to yield gene trap vectors for all three forward frames.

sgRNA rx3_T1 (AGCAGAGCGCGCAAAGAACC[AGG], PAM in brackets) and rx3_T2 (AGCGCGCAAAGAACCAGGCA[GGG], PAM in brackets) were designed using CCTop and cloned and transcribed as described previously (Stemmer et al., 2015). The saGFP-OPT-MCS+2 cassette was inserted into the first intron of *Rx3* by non-homologous end-joining *via* microinjection into the cytoplasm of one cell stage medaka zygotes. Injection mix contained 15 ng/µl of each sgRNA *Rx3_T1, Rx3_T2* and *GFP_T1*, 150 ng/µl Cas9 mRNA and 10 ng/µl pGGD(saGFP-OPT-MCS+2) in nuclease-free water. Embryos were raised and maintained at 28°C in 1× Embryo Rearing Medium (ERM, 17 mM NaCl, 40 mM KCl, 0.27 mM CaCl_2_, 0.66 mM MgSO_4_, 17 mM HEPES) and screened for ocular GFP expression 1 day post fertilization on a Nikon SMZ18. Genotyping-PCR was performed with Q5^®^ High-Fidelity DNA Polymerase (NEB) with 98°C initial denaturation for 2 min, followed by 30 cycles of: 98°C denaturation for 20 s, 66°C annealing for 30 s, 72°C extension for 25 s. Primers used to amplify 5’ integration: Rx3_F 5’-TCCTTTTTAGACAAATGTGGCTCC, GFP_R 5’-GCTCGACCAGGATGGGCA; 3’ integration: pDest_F 5’-ATTACCGCCTTTGAGTGAGC, Rx3_R 5’-GACAGGTATCCGGTAAGCGG. Following gel electrophoresis, amplicons were gel purified (InnuPrep, AnalyticJena), sequenced (Eurofins Genomics) and analyzed using Geneious R8.1 (Biomatters).

### Injection in zebrafish embryos

Rx3::H2B-GFP DNA (10 ng/μl) was co-injected with Meganuclease (I-SceI) and mCherry mRNA (10 ng/μl) into the cytoplasm of one cell stage zebrafish embryos as previously described (Thermes et al., 2002). Only morphologically intact and brightest mCherry expressing embryos were used for aggregate formation.

### Generation of organoids

Blastula stage (6 hpf) embryos (Iwamatsu, 2004) were collected, dechorionated using hatching enzyme and washed in Embryo Rearing Media (ERM, 17 mM NaCl, 40 mM KCl, 0.27 mM CaCl_2_, 0.66 mM MgSO_4_, 17 mM HEPES). Cell mass was separated from yolk, washed 3 times with sterile PBS and dissociated by gentle pipetting with 200 μl pipet tip. Cell suspension was pelleted (180 x g for 3 min) and re-suspended in differentiation media: GMEM (Glasgow’ s Minimal Essential Medium, Gibco), 5% KSR (knockout serum replacement, Gibco), 0.1 mM non-essential amino acids, sodium pyruvate, 0.1 mM β-mercaptoethanol, 50 U/ml penicillin-streptomycin to desired cell density, i.e. regular aggregates 10–20 cells/µl and small aggregates 5–8 cells/µl. Cell suspension (100 μl per well-per single aggregate) was transferred to low binding 96-well plate (Nunclon Sphera U-Shaped Bottom Microplate, Thermo Fisher Scientific) and centrifugated (for 3 min at 180 × g) to speed up cell aggregation. The following day (day 1) aggregates were washed with differentiation media, transferred to fresh wells and Matrigel (Corning, 356238) was added to the media to a final concentration of 2%. From day 2 onwards organoids were kept in DMEM/F12 supplemented with 5% FBS, 5% FEE (fish embryonic extract) (https://zfin.org/zf_info/zfbook/chapt6.html), 20 mM HEPES pH=7.4, N2 supplement (Gibco) and 50 U/ml penicillin-streptomycin. For zebrafish cell aggregation, blastula-stage embryos were processed according to the same protocol and re-suspended in Leibowitz’ s L-15 media supplemented with 5% KSR and penicillin-streptomycin. Matrigel (Corning, 356238) was added 2 hpa.

### Immunohistochemistry

Immunohistochemistry was performed as previously described (Inoue and Wittbrodt, 2011) with slight modifications. Embryos and organoids were processed as whole-mounts, fixed in 4% PFA overnight at 4°C, washed in PTW (PBS with 0.05% Tween20), heated in 50 mM Tris-HCl at 70°C for 15 min, permeabilized 15 min in acetone at –20°C and blocked in 10% BSA in PTW for 1 h. Samples were incubated with primary antibody (1:200) overnight (for Otx2 antibody incubation was prolonged to three days) at 4°C. The following antibodies were used: rabbit anti-Rx2 (Reinhardt *et al*., 2015), rabbit anti-Lhx2 (GeneTex, GTX129241), rabbit anti-β-catenin (Abcam, Ab6302), mouse anti-acetylated tubulin (Sigma, T7451), rabbit anti-Sox2 (GeneTex, GTX124477), chicken anti-GFP (Life Technologies, A10262), goat anti-Otx2 (R&D, AF1979), rabbit anti-Prox1 (Millipore, AB5475) and mouse anti-HuC/D (Life Technologies, A21271). Samples were washed 6 times 10 min in PTW, incubated with secondary antibody (1:750) (Invitrogen) with DAPI (1:500) overnight at 4°C, washed 5 times 10 min in PTW, mounted in 1% low melting agarose in PTW and imaged with Sp8 Leica confocal microscope.

### Imaging

Time lapse imaging of aggregation and *Rx3* expression profiles in organoids was performed on the ACQUIFER Imaging Machine (ACQUIFER Imaging GmbH, Heidelberg, Germany) (Pandey et al., 2019). Aggregates and dechorionated embryos were loaded into 96-well plates and placed in the plate holder of the ACQUIFER machine at 26°C for medaka and 28°C for zebrafish. For each well plate, a set of 10 z-slices (75-µm step size) was acquired in the brightfield (50% LED intensity, 50 ms exposure time) and 470 nm fluorescence (50% LED excitation source, FITC channel, 200 ms exposure time) channels with a 4× NA 0.13 objective (Nikon, Düsseldorf, Germany). Imaging was performed over 20 h of organoid development with 30 min intervals.

3D imaging of fixed *Atoh7::EGFP*-derived organoids was performed on multiview selective-plane illumination MuVi SPIM Multiview light-sheet microscope (Luxendo Light-sheet, Bruker Corporation) (Krzic et al., 2012). The organoids were mounted in 1% low-melting agarose (StarPure Low Melt Agarose, cat. no. N3103-0100, StarLab GmbH) inside a FEP tube (Karl Schupp AG) fixed on glass capillaries. Four volumes (two cameras and two rotation angles) were acquired with the 25× detection setup (Caroti et al., 2018), 1-µm z step size for 3 channels: GFP (Atoh7) 488 nm excitation laser at 40% intensity, 525/50 nm emission filter, 600 ms exposure time; RFP (β-catenin) 561 nm excitation laser at 40% intensity, 607/70 nm emission filter, 600 ms exposure time; far Red (HuC/D) 642 nm excitation laser at 50% intensity, 579/40 nm emission filter, 1000 ms exposure time. The volumes were fused with Luxendo Image fusion software. Fused volumes were visualized by 3D rendering with Amira 6.2 Main software.

Live imaging of *Rx3::H2B-GFP*-derived organoids was performed on the 16× detection MuVi SPIM Multiview light-sheet microscope (Luxendo Light-sheet, Bruker Corporation). To assure normal development of organoids, the FEP tube was partially filled with 2% low-melting agarose. After solidification of agarose the tube was further filled with differentiation media and an organoid was scooped inside the tube. The tube was positioned vertically such that the organoid fell onto the solid support of agarose. Two volumes with 1.6-µm z step size for 2 channels, i.e. GFP 488 nm excitation laser at 10% intensity, 525/50 nm emission filter, 50 ms exposure time and quasi bright field 561 nm excitation laser at 5% intensity, 568LP nm emission filter, 100 ms exposure time, were acquired every 15 min.

Live imaging of *Rx3::H2B-GFP*-derived organoids (Video 5 and 6) was performed with Sp8 confocal microscope (Leica). Organoids were imaged from day 1 to day 2 at room temperature directly in low binding 96-well plate (Nunclon Sphera U-Shaped Bottom Microplate, Thermo Fisher Scientific). The volume of the organoids was acquired with 2.24 µm z step size using 488 nm excitation laser at 10% intensity in 30 min intervals.

Fixed organoid and embryonic samples were imaged with Sp8 confocal microscope (Leica). Gross morphology of embryos and organoids was assessed by Nikon SMZ18 and Leica DMi8 microscope.

### Quantitative analysis

To analyze the dynamics of retinal cell fate acquisition, for each time point, a maximum projection for fluorescence and a single focused bright-field image were calculated using Fiji distribution of ImageJ (Schindelin et al., 2012) and hyper stack plugin for ACQUIFER machine (https://doi.org/10.5281/zenodo.3368134). The onset of GFP expression was determined by detection of maximum in first derivative of GFP expression. For better representation, the sum of fluorescence intensity for each time point was normalized from 0 to 1.

Level of organization in organoids derived from *Rx3*-mutant blastulae was performed on images acquired with Leica DMi8 Microscope. Two step automatic thresholding (Minimum, Fiji (REF)) resulted in two regions, the first corresponding to the whole area (A1) of an organoid and the second to the unorganized (dark) region (A2) within it. Level of organization was determined as 1-A2/A1 and grouped in two phenotypes by k-Means clustering. A similar approach was used for analysis of gene expression in regular and small aggregates. Automatic thresholding on bright channel was used to determine area of an organoid and on fluorescence image to identify areas corresponding to Rx3 or Rx2 expression.

To analyze cell migration behavior in organoids of day 2, time lapse volumes acquired with SPIM were registered with ElastixWrapper for Fiji (Christian Tischer, 2019; Klein et al., 2010) using Euler transformation, 1000 iterations, full data points. The cells within organoids were tracked with TrackMate (Tinevez et al., 2017), using simple linear tracker. The analysis of tracks’ directionality was performed with custom-written Matlab script. The MSD analysis including velocity autocorrelation was performed with Matlab tool for analysis of particle trajectories (Tinevez and Herbert, 2020). For demonstration purposes, we chose a few representative tracks based on total displacement (total distance traveled) and duration (time required for travel).

Statistical analysis and plots were prepared with Jupyter notebook (Perez and Granger, 2007), using statannot package (https://github.com/webermarcolivier/statannot) to compute statistical tests and add statistical annotations. For all figures, ns 0.05<p<1, ∗ 0.01<p<0.05, ∗∗ 0.001<p<0.01, ∗∗∗ 0.0001<p<0.001, ∗∗∗∗ p<0.0001. Figures were assembled with Adobe Illustrator CS6.

## Acknowledgements

The authors thank L. Centanin, E. Tsingos, N. Sokolova and Z. Kozmik for valuable discussions and comments on the manuscript. We are grateful to I. Thomas, A. Sarvari, M. Pandya, C. Schlagheck and N. Priya for the help at the initial stage of the project. We thank Darius Balciunas for providing the GBT-RP2 plasmid containing the OPT sequence. V.W. was supported by the German Research Foundation research fellowship WE 6221/2-1. This work was supported by grants of the Excellence Cluster “3D Matter Made to Order” (3DMM2O) funded through the German Excellence Strategy via Deutsche Forschungsgemeinschaft (DFG), by the Carl Zeiss Foundation and by the ERC Synergy Grant IndiGene (Number 810172) to JW.

## Video legends

**Video 1. Aggregation and compaction of blastula-derived primary embryonic stem cells**. Time-lapse imaging of medaka-derived blastula cells going through the process of aggregation and compaction. Imaging was performed with 30 min intervals. Scale bar: 100 μm.

**Video 2. Acquisition of retinal fate within medaka blastula-derived aggregates**. Time-lapse imaging of *Rx3::H2B-GFP* medaka-derived blastula cells going through the process of aggregation, compaction and acquisition of retinal fate (GFP expression). Imaging was performed with 30 min intervals. Scale bar: 100 μm.

**Video 3. Acquisition of retinal fate within zebrafish blastula-derived aggregates**. Time-lapse imaging of *Rx3::H2B-GFP* zebrafish-derived blastula cells going through the process of aggregation, compaction and acquisition of retinal fate (GFP expression). Imaging was performed with 30 min intervals. Scale bar: 100 μm.

**Video 4. Optic vesicle formation within blastula-derived retinal organoid**. Time-lapse imaging of optic vesicle-like structure evagination in medaka *Rx3::H2B-GFP* retinal organoids. Maximal projection of quasi bright-field, GFP and tracks of exemplary cells. Imaging was performed with 15 min intervals. Scale bar: 100 μm.

**Video 5. Example of organoids forming single and multiple optic vesicle-like structures**. Time-lapse imaging of optic vesicle-like structure evagination in medaka *Rx3::H2B-GFP* retinal organoids. Maximal projection of GFP and tracks of exemplary cells. Imaging was performed with 15 min intervals. Scale bar: 100 μm.

**Video 6. Extracellular matrix controls formation of optic vesicle neuroepithelium**. Time-lapse imaging of optic vesicle-like structure evagination in medaka *Rx3::H2B-GFP* retinal organoids in presence or absence of Matrigel (extracellular matrix component) shown as maximal projection of GFP and brightfield. Imaging was performed with 30 min intervals. Scale bar: 100 μm.

**Video 7. Organoids show complex morphology with the onset of retinal differentiation**. 3D rendering of zebrafish *Atoh7::EGFP* day 3 organoid stained with anti-GFP (green), anti-β-catenin (cyan) and anti HuC/D (magenta) antibodies.

## Supplementary Figures

**Figure S1.**
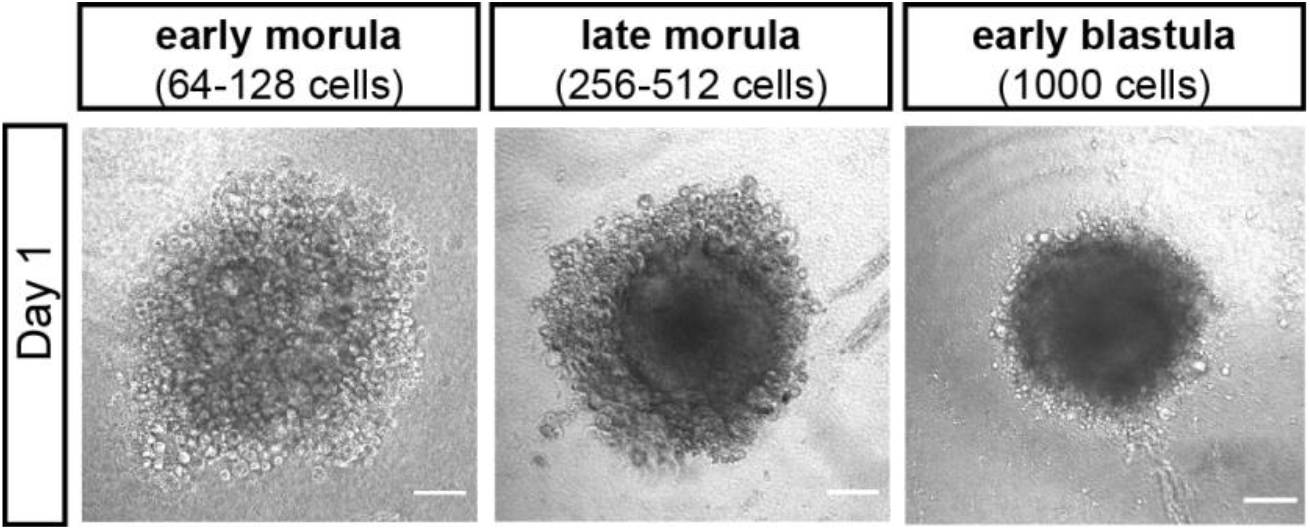
Impact of time point of cell extraction on aggregation efficiency. Bright-field images of day 1 aggregates derived from early morula (64–128 cells), late morula (256–512 cells) and early blastula (1,000 cells). Scale bars: 100 μm.

**Figure S2.**
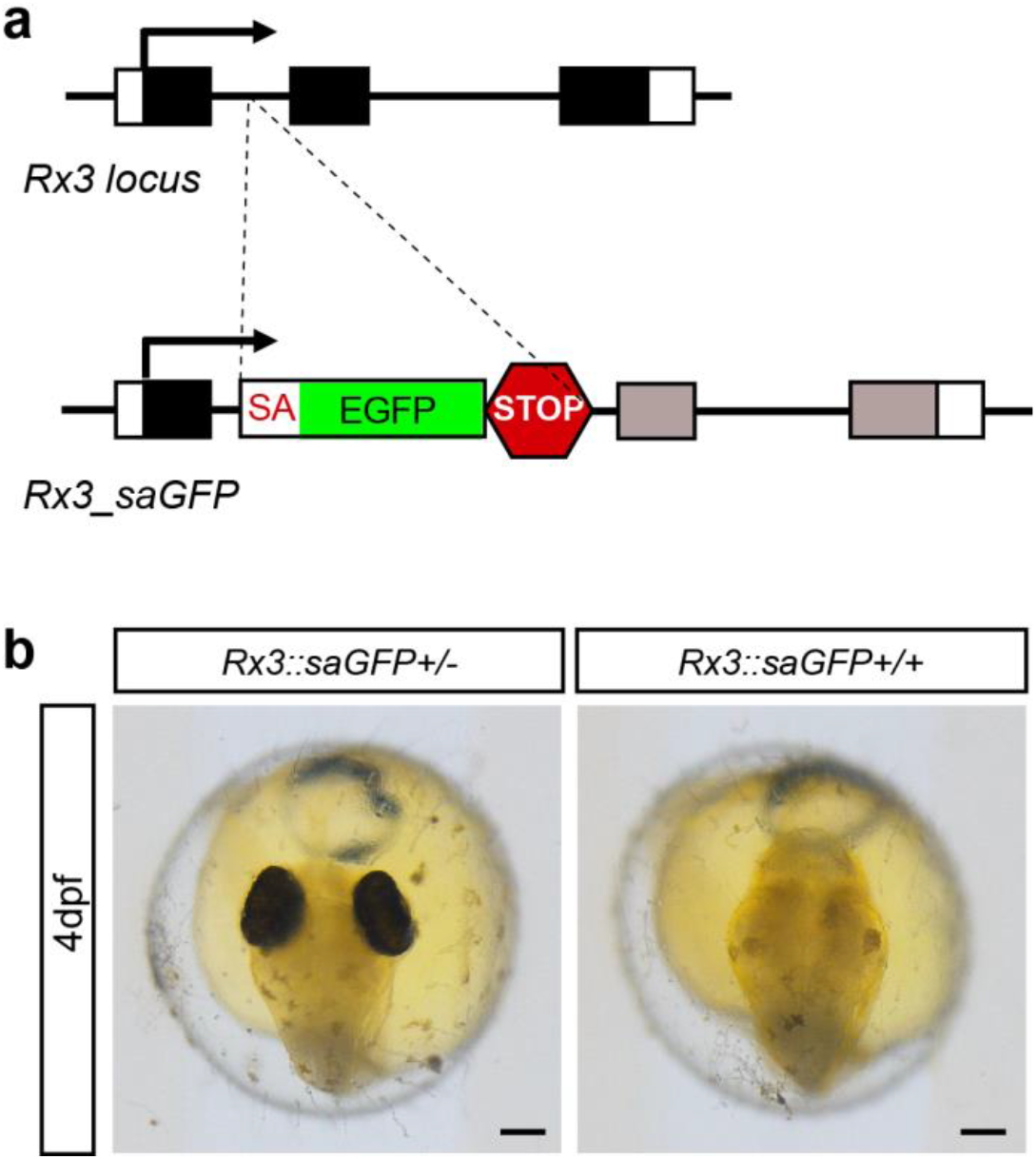
Generation and characterization of *Rx3KO* (*Rx3::saGFP*) line. **(a)** Schematic representation of *Rx3* locus with integrated saGFP-OPT cassette. An open reading frame-adjusted gene trap cassette comprising a splice acceptor and a *GFP* sequence (saGFP) followed by a polyA and strong terminator sequence derived from the ocean pout (OPT; STOP) were inserted in the *Rx3* locus. **(b)** Bright-field images of fish embryos 4 dpf showing the phenotypical consequence of cassette integration in the *Rx3* locus. dpf, days post fertilization. Scale bars: 100 μm.

**Figure S3.**
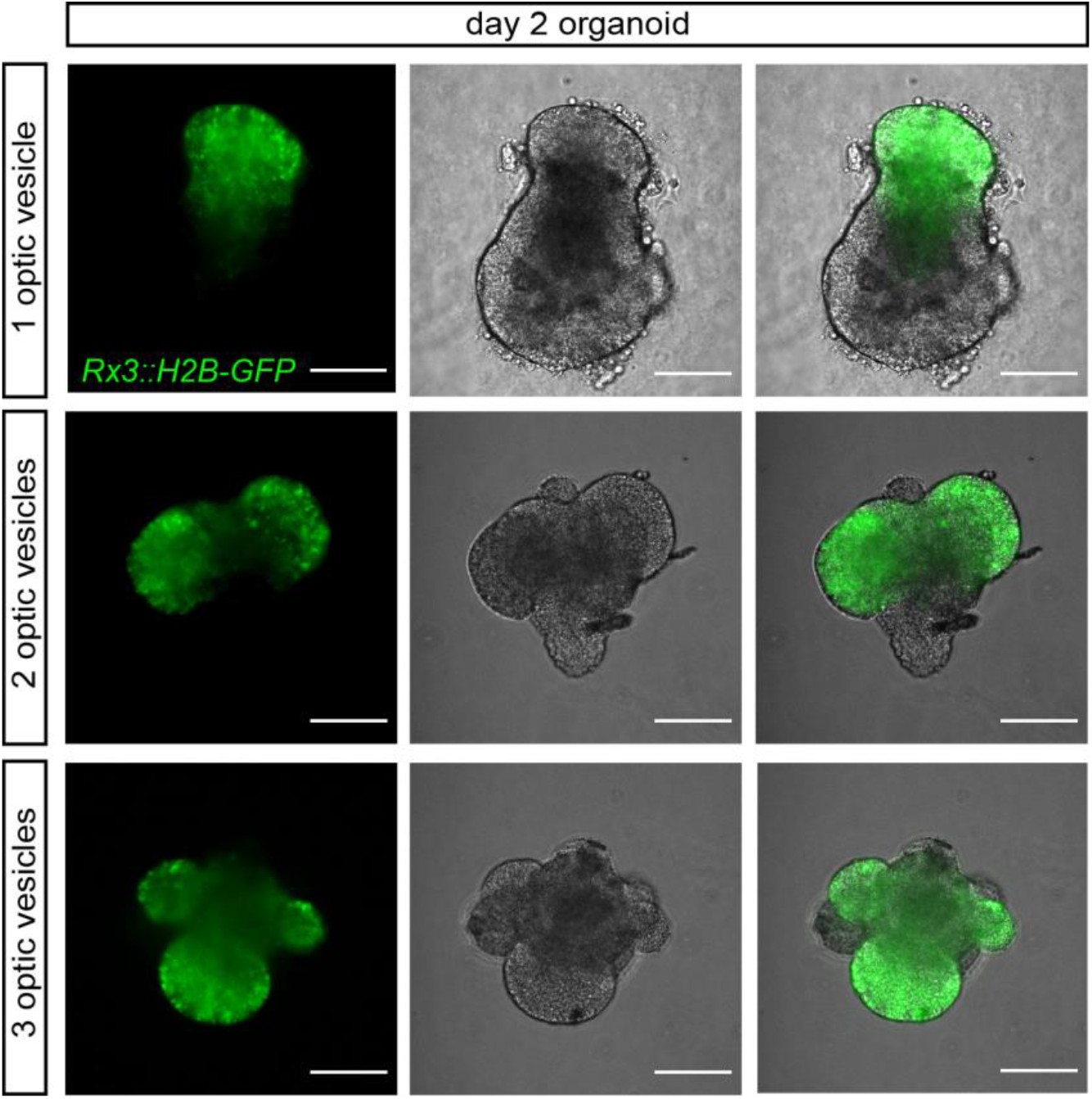
Examples of small organoids forming one, two and three optic vesicle-like structures. Fluorescence, bright-field images and their overlay of retinal organoids generated from *Rx3::H2B-GFP* reporter line formed by aggregation of 500–800 primary embryonic stem cells stained with anti-GFP antibody. Scale bars: 100 μm.

## References

Acampora D, Mazan S, Lallemand Y, Avantaggiato V, Maury M, Simeone A, Brûlet P. 1995. Forebrain and midbrain regions are deleted in Otx2-/-mutants due to a defective anterior neuroectoderm specification during gastrulation. Dev Camb Engl 121:3279– 3290.

Azizi A, Herrmann A, Wan Y, Buse SJ, Keller PJ, Goldstein RE, Harris WA. 2020. Nuclear crowding and nonlinear diffusion during interkinetic nuclear migration in the zebrafish retina. eLife 9:e58635. doi:10.7554/eLife.58635

Beccari L, Moris N, Girgin M, Turner DA, Baillie-Johnson P, Cossy A-C, Lutolf MP, Duboule D, Arias AM. 2018. Multi-axial self-organization properties of mouse embryonic stem cells into gastruloids. Nature 562:272–276. doi:10.1038/s41586-018-0578-0

Bovolenta P, Mallamaci A, Briata P, Corte G, Boncinelli E. 1997. Implication of OTX2 in Pigment Epithelium Determination and Neural Retina Differentiation. J Neurosci 17:4243–4252. doi:10.1523/JNEUROSCI.17-11-04243.1997

Brink SC van den, Baillie-Johnson P, Balayo T, Hadjantonakis A-K, Nowotschin S, Turner DA, Arias AM. 2014. Symmetry breaking, germ layer specification and axial organisation in aggregates of mouse embryonic stem cells. Development 141:4231– 4242. doi:10.1242/dev.113001

Brooks MJ, Chen HY, Kelley RA, Mondal AK, Nagashima K, De Val N, Li T, Chaitankar V, Swaroop A. 2019. Improved Retinal Organoid Differentiation by Modulating Signaling Pathways Revealed by Comparative Transcriptome Analyses with Development In Vivo. Stem Cell Rep 13:891–905. doi:10.1016/j.stemcr.2019.09.009

Brown NL, Kanekar S, Vetter ML, Tucker PK, Gemza DL, Glaser T. 1998. Math5 encodes a murine basic helix-loop-helix transcription factor expressed during early stages of retinal neurogenesis. Dev Camb Engl 125:4821–4833.

Caroti F, González Avalos E, Noeske V, González Avalos P, Kromm D, Wosch M, Schütz L, Hufnagel L, Lemke S. 2018. Decoupling from yolk sac is required for extraembryonic tissue spreading in the scuttle fly Megaselia abdita. eLife 7. doi:10.7554/eLife.34616

Centanin L, Hoeckendorf B, Wittbrodt J. 2011. Fate Restriction and Multipotency in Retinal Stem Cells. Cell Stem Cell 9:553–562. doi:10.1016/j.stem.2011.11.004

Chauhan B, Plageman T, Lou M, Lang R. 2015. Chapter Eleven - Epithelial Morphogenesis: The Mouse Eye as a Model System1 In: Trainor PA, editor. Current Topics in Developmental Biology, Neural Crest and Placodes. Academic Press. pp. 375–399. doi:10.1016/bs.ctdb.2014.11.011

Chow RL, Lang RA. 2001. Early Eye Development in Vertebrates. Annu Rev Cell Dev Biol 17:255–296. doi:10.1146/annurev.cellbio.17.1.255

Christian Tischer. 2019. ElastixWrapper: Fiji plugin for 3D image registration with elastix. Zenodo. doi:10.5281/zenodo.2602549

Chuang JC, Mathers PH, Raymond PA. 1999. Expression of three Rx homeobox genes in embryonic and adult zebrafish. Mech Dev 84:195–198. doi:10.1016/s0925-4773(99)00077-5

Clark KJ, Balciunas D, Pogoda H-M, Ding Y, Westcot SE, Bedell VM, Greenwood TM, Urban MD, Skuster KJ, Petzold AM, Ni J, Nielsen A, Patowary A, Scaria V, Sivasubbu S, Xu X, Hammerschmidt M, Ekker SC. 2011. in vivo protein trapping produces a functional expression codex of the vertebrate proteome. Nat Methods 8:506–515. doi:10.1038/nmeth.1606

Del Bene F, Ettwiller L, Skowronska-Krawczyk D, Baier H, Matter J-M, Birney E, Wittbrodt J. 2007. In vivo validation of a computationally predicted conserved Ath5 target gene set. PLoS Genet 3:1661–1671. doi:10.1371/journal.pgen.0030159

Deschet K, Bourrat F, Ristoratore F, Chourrout D, Joly J-S. 1999. Expression of the medaka (Oryzias latipes) Ol-Rx3 paired-like gene in two diencephalic derivatives, the eye and the hypothalamus. Mech Dev 83:179–182. doi:10.1016/S0925-4773(99)00037-4

Dyer MA, Livesey FJ, Cepko CL, Oliver G. 2003. Prox1 function controls progenitor cell proliferation and horizontal cell genesis in the mammalian retina. Nat Genet 34:53–58. doi:10.1038/ng1144

Eiraku M, Takata N, Ishibashi H, Kawada M, Sakakura E, Okuda S, Sekiguchi K, Adachi T, Sasai Y. 2011. Self-organizing optic-cup morphogenesis in three-dimensional culture. Nature 472:51–56. doi:10.1038/nature09941

Fligor CM, Langer KB, Sridhar A, Ren Y, Shields PK, Edler MC, Ohlemacher SK, Sluch VM, Zack DJ, Zhang C, Suter DM, Meyer JS. 2018. Three-Dimensional Retinal Organoids Facilitate the Investigation of Retinal Ganglion Cell Development, Organization and Neurite Outgrowth from Human Pluripotent Stem Cells. Sci Rep 8:14520. doi:10.1038/s41598-018-32871-8

Fossat N, Le Greneur C, Béby F, Vincent S, Godement P, Chatelain G, Lamonerie T. 2007. A new GFP-tagged line reveals unexpected Otx2 protein localization in retinal photoreceptors. BMC Dev Biol 7:122. doi:10.1186/1471-213X-7-122

Fuhrmann S. 2010. Eye Morphogenesis and Patterning of the Optic Vesicle. Curr Top Dev Biol 93:61–84. doi:10.1016/B978-0-12-385044-7.00003-5

Fulton T, Trivedi V, Attardi A, Anlas K, Dingare C, Arias AM, Steventon B. 2020. Axis Specification in Zebrafish Is Robust to Cell Mixing and Reveals a Regulation of Pattern Formation by Morphogenesis. Curr Biol 30:3063–3064. doi:10.1016/j.cub.2020.07.022

Gao M-L, Lei X-L, Han F, He K-W, Jin S-Q, Zhang Y-Y, Jin Z-B. 2020. Patient-Specific Retinal Organoids Recapitulate Disease Features of Late-Onset Retinitis Pigmentosa. Front Cell Dev Biol 8. doi:10.3389/fcell.2020.00128

Glubrecht DD, Kim J-H, Russell L, Bamforth JS, Godbout R. 2009. Differential CRX and OTX2 expression in human retina and retinoblastoma. J Neurochem 111:250–263. doi:10.1111/j.1471-4159.2009.06322.x

Good PJ. 1995. A conserved family of elav-like genes in vertebrates. Proc Natl Acad Sci U S A 92:4557–4561.

Gutierrez-Triana JA, Tavhelidse T, Thumberger T, Thomas I, Wittbrodt B, Kellner T, Anlas K, Tsingos E, Wittbrodt J. 2018. Efficient single-copy HDR by 5’ modified long dsDNA donors. eLife 7:e39468. doi:10.7554/eLife.39468

Hatakeyama J, Tomita K, Inoue T, Kageyama R. 2001. Roles of homeobox and bHLH genes in specification of a retinal cell type. Development 128:1313–1322.

Hemmati-Brivanlou A, Melton D. 1997. Vertebrate embryonic cells will become nerve cells unless told otherwise. Cell 88:13–17. doi:10.1016/s0092-8674(00)81853-x

Hirashima M, Kobayashi T, Uchikawa M, Kondoh H, Araki M. 2008. Anteroventrally localized activity in the optic vesicle plays a crucial role in the optic development. Dev Biol 317:620–631. doi:10.1016/j.ydbio.2008.03.010

Ho SY, Goh CWP, Gan JY, Lee YS, Lam MKK, Hong N, Hong Y, Chan WK, Shu-Chien AC. 2014. Derivation and Long-Term Culture of an Embryonic Stem Cell-Like Line from Zebrafish Blastomeres Under Feeder-Free Condition. Zebrafish 11:407. doi:10.1089/zeb.2013.0879

Hong Y, Winkler C, Schartl M. 1998. Production of medakafish chimeras from a stable embryonic stem cell line. Proc Natl Acad Sci 95:3679–3684. doi:10.1073/pnas.95.7.3679

Hong Y, Winkler C, Schartl M. 1996. Pluripotency and differentiation of embryonic stem cell lines from the medakafish (Oryzias latipes). Mech Dev 60:33–44. doi:10.1016/s0925-4773(96)00596-5

Inoue D, Wittbrodt J. 2011. One for all--a highly efficient and versatile method for fluorescent immunostaining in fish embryos. PloS One 6:e19713. doi:10.1371/journal.pone.0019713

Ivanovitch K, Cavodeassi F, Wilson SW. 2013. Precocious acquisition of neuroepithelial character in the eye field underlies the onset of eye morphogenesis. Dev Cell 27:293– 305. doi:10.1016/j.devcel.2013.09.023

Iwamatsu T. 2004. Stages of normal development in the medaka Oryzias latipes. Mech Dev, Medaka 121:605–618. doi:10.1016/j.mod.2004.03.012

Kay JN, Finger-Baier KC, Roeser T, Staub W, Baier H. 2001. Retinal Ganglion Cell Genesis Requires lakritz, a Zebrafish atonal Homolog. Neuron 30:725–736. doi:10.1016/S0896-6273(01)00312-9

Kim C-H, Ueshima E, Muraoka O, Tanaka H, Yeo S-Y, Huh T-L, Miki N. 1996. Zebra?sh elav/HuC homologue as a very early neuronal marker. Neurosci Lett 4.

Kimmel CB, Ballard WW, Kimmel SR, Ullmann B, Schilling TF. 1995. Stages of embryonic development of the zebrafish. Dev Dyn Off Publ Am Assoc Anat 203:253–310. doi:10.1002/aja.1002030302

Kirchmaier S, Lust K, Wittbrodt J. 2013. Golden GATEway Cloning – A Combinatorial Approach to Generate Fusion and Recombination Constructs. PLOS ONE 8:e76117. doi:10.1371/journal.pone.0076117

Klein S, Staring M, Murphy K, Viergever MA, Pluim JPW. 2010. elastix: a toolbox for intensity-based medical image registration. IEEE Trans Med Imaging 29:196–205. doi:10.1109/TMI.2009.2035616

Koike C, Nishida A, Ueno S, Saito H, Sanuki R, Sato S, Furukawa A, Aizawa S, Matsuo I, Suzuki N, Kondo M, Furukawa T. 2007. Functional Roles of Otx2 Transcription Factor in Postnatal Mouse Retinal Development. Mol Cell Biol 27:8318–8329. doi:10.1128/MCB.01209-07

Kruczek K, Swaroop A. 2020. Pluripotent stem cell-derived retinal organoids for disease modeling and development of therapies. STEM CELLS 38:1206–1215. doi:https://doi.org/10.1002/stem.3239

Krzic U, Gunther S, Saunders TE, Streichan SJ, Hufnagel L. 2012. Multiview light-sheet microscope for rapid in toto imaging. Nat Methods 9:730–733. doi:10.1038/nmeth.2064

Kuwahara A, Ozone C, Nakano T, Saito K, Eiraku M, Sasai Y. 2015. Generation of a ciliary margin-like stem cell niche from self-organizing human retinal tissue. Nat Commun 6:6286. doi:10.1038/ncomms7286

Lancaster MA, Knoblich JA. 2014. Generation of Cerebral Organoids from Human Pluripotent Stem Cells. Nat Protoc 9:2329–2340. doi:10.1038/nprot.2014.158

Levine AJ, Brivanlou AH. 2007. Proposal of a model of mammalian neural induction. Dev Biol 308:247–256. doi:10.1016/j.ydbio.2007.05.036

Li H, Tierney C, Wen L, Wu JY, Rao Y. 1997. A single morphogenetic field gives rise to two retina primordia under the influence of the prechordal plate. Dev Camb Engl 124:603– 615.

Link BA, Fadool JM, Malicki J, Dowling JE. 2000. The zebrafish young mutation acts non-cell-autonomously to uncouple differentiation from specification for all retinal cells. Development 127:2177–2188.

Loosli F, Staub W, Finger-Baier KC, Ober EA, Verkade H, Wittbrodt J, Baier H. 2003. Loss of eyes in zebrafish caused by mutation of chokh/rx3. EMBO Rep 4:894– 899. doi:10.1038/sj.embor.embor919

Loosli F, Winkler S, Burgtorf C, Wurmbach E, Ansorge W, Henrich T, Grabher C, Arendt D, Carl M, Krone A, Grzebisz E, Wittbrodt J. 2001. Medaka eyeless is the key factor linking retinal determination and eye growth. Development 128:4035–4044.

Loosli F, Winkler S, Wittbrodt J. 1999. Six3 overexpression initiates the formation of ectopic retina. Genes Dev 13:649–654.

Martinez-Morales J-R, Cavodeassi F, Bovolenta P. 2017. Coordinated Morphogenetic Mechanisms Shape the Vertebrate Eye. Front Neurosci 11. doi:10.3389/fnins.2017.00721

Mathers PH, Grinberg A, Mahon KA, Jamrich M. 1997. The Rx homeobox gene is essential for vertebrate eye development. Nature 387:603–607. doi:10.1038/42475

Meinhardt H. 2012. Turing’ s theory of morphogenesis of 1952 and the subsequent discovery of the crucial role of local self-enhancement and long-range inhibition. Interface Focus 2:407–416. doi:10.1098/rsfs.2011.0097

Moris N, Anlas K, van den Brink SC, Alemany A, Schröder J, Ghimire S, Balayo T, van Oudenaarden A, Martinez Arias A. 2020. An in vitro model of early anteroposterior organization during human development. Nature 582:410–415. doi:10.1038/s41586-020-2383-9

Nakano T, Ando S, Takata N, Kawada M, Muguruma K, Sekiguchi K, Saito K, Yonemura S, Eiraku M, Sasai Y. 2012. Self-formation of optic cups and storable stratified neural retina from human ESCs. Cell Stem Cell 10:771–785. doi:10.1016/j.stem.2012.05.009

Nishida A, Furukawa A, Koike C, Tano Y, Aizawa S, Matsuo I, Furukawa T. 2003. Otx2 homeobox gene controls retinal photoreceptor cell fate and pineal gland development. Nat Neurosci 6:1255–1263. doi:10.1038/nn1155

Pandey G, Westhoff JH, Schaefer F, Gehrig J. 2019. A Smart Imaging Workflow for Organ-Specific Screening in a Cystic Kidney Zebrafish Disease Model. Int J Mol Sci 20:1290. doi:10.3390/ijms20061290

Park HC, Kim CH, Bae YK, Yeo SY, Kim SH, Hong SK, Shin J, Yoo KW, Hibi M, Hirano T, Miki N, Chitnis AB, Huh TL. 2000. Analysis of upstream elements in the HuC promoter leads to the establishment of transgenic zebrafish with fluorescent neurons. Dev Biol 227:279–293. doi:10.1006/dbio.2000.9898

Peng L, Zhou Y, Xu W, Jiang M, Li H, Long M, Liu W, Liu J, Zhao X, Xiao Y. 2019. Generation of Stable Induced Pluripotent Stem-like Cells from Adult Zebra Fish Fibroblasts. Int J Biol Sci 15:2340–2349. doi:10.7150/ijbs.34010

Perez F, Granger BE. 2007. IPython: A System for Interactive Scientific Computing. Comput Sci Eng 9:21–29. doi:10.1109/MCSE.2007.53

Pevny L, Placzek M. 2005. SOX genes and neural progenitor identity. Curr Opin Neurobiol 15:7–13. doi:10.1016/j.conb.2005.01.016

Poggi L, Vitorino M, Masai I, Harris WA. 2005. Influences on neural lineage and mode of division in the zebrafish retina in vivo. J Cell Biol 171:991–999. doi:10.1083/jcb.200509098

Prykhozhij SV, Berman JN. 2018. Zebrafish knock-ins swim into the mainstream. Dis Model Mech 11. doi:10.1242/dmm.037515

Rembold M, Loosli F, Adams RJ, Wittbrodt J. 2006. Individual Cell Migration Serves as the Driving Force for Optic Vesicle Evagination. Science 313:1130–1134. doi:10.1126/science.1127144

Robles V, Martĺ M, Izpisua Belmonte JC. 2011. Study of pluripotency markers in zebrafish embryos and transient embryonic stem cell cultures. Zebrafish 8:57–63. doi:10.1089/zeb.2010.0684

Schauer A, Pinheiro D, Hauschild R, Heisenberg C-P. 2020. Zebrafish embryonic explants undergo genetically encoded self-assembly. eLife 9:e55190. doi:10.7554/eLife.55190

Schindelin J, Arganda-Carreras I, Frise E, Kaynig V, Longair M, Pietzsch T, Preibisch S, Rueden C, Saalfeld S, Schmid B, Tinevez J-Y, White DJ, Hartenstein V, Eliceiri K, Tomancak P, Cardona A. 2012. Fiji: an open-source platform for biological-image analysis. Nat Methods 9:676–682. doi:10.1038/nmeth.2019

Simeone A, Acampora D, Mallamaci A, Stornaiuolo A, D’ Apice MR, Nigro V, Boncinelli E. 1993. A vertebrate gene related to orthodenticle contains a homeodomain of the bicoid class and demarcates anterior neuroectoderm in the gastrulating mouse embryo. EMBO J 12:2735–2747.

Stemmer M, Thumberger T, Keyer M del S, Wittbrodt J, Mateo JL. 2015. CCTop: An Intuitive, Flexible and Reliable CRISPR/Cas9 Target Prediction Tool. PLOS ONE 10:e0124633. doi:10.1371/journal.pone.0124633

Stigloher C, Ninkovic J, Laplante M, Geling A, Tannhäuser B, Topp S, Kikuta H, Becker TS, Houart C, Bally-Cuif L. 2006. Segregation of telencephalic and eye-field identities inside the zebrafish forebrain territory is controlled by Rx3. Development 133:2925– 2935. doi:10.1242/dev.02450

Thermes V, Grabher C, Ristoratore F, Bourrat F, Choulika A, Wittbrodt J, Joly J-S. 2002. I-SceI meganuclease mediates highly efficient transgenesis in fish. Mech Dev 118:91– 98. doi:10.1016/S0925-4773(02)00218-6

Tian E, Kimura C, Takeda N, Aizawa S, Matsuo I. 2002. Otx2 is required to respond to signals from anterior neural ridge for forebrain specification. Dev Biol 242:204–223. doi:10.1006/dbio.2001.0531

Tinevez J-Y, Herbert S. 2020. The NEMO Dots Assembly: Single-Particle Tracking and Analysis In: Miura K, Sladoje N, editors. Bioimage Data Analysis Workflows, Learning Materials in Biosciences. Cham: Springer International Publishing. pp. 67– 96. doi:10.1007/978-3-030-22386-1_4

Tinevez J-Y, Perry N, Schindelin J, Hoopes GM, Reynolds GD, Laplantine E, Bednarek SY, Shorte SL, Eliceiri KW. 2017. TrackMate: An open and extensible platform for single-particle tracking. Methods San Diego Calif 115:80–90. doi:10.1016/j.ymeth.2016.09.016

Tropepe V, Hitoshi S, Sirard C, Mak TW, Rossant J, van der Kooy D. 2001. Direct neural fate specification from embryonic stem cells: a primitive mammalian neural stem cell stage acquired through a default mechanism. Neuron 30:65–78. doi:10.1016/s0896-6273(01)00263-x

Turing AM. 1952. The chemical basis of morphogenesis. Philos Trans R Soc Lond B Biol Sci 237:37–72. doi:10.1098/rstb.1952.0012

Turner DA, Girgin M, Alonso-Crisostomo L, Trivedi V, Baillie-Johnson P, Glodowski CR, Hayward PC, Collignon J, Gustavsen C, Serup P, Steventon B, Lutolf MP, Arias AM. 2017. Anteroposterior polarity and elongation in the absence of extra-embryonic tissues and of spatially localised signalling in gastruloids: mammalian embryonic organoids. Development 144:3894–3906. doi:10.1242/dev.150391

Turner DA, Trott J, Hayward P, Rué P, Arias AM. 2014. An interplay between extracellular signalling and the dynamics of the exit from pluripotency drives cell fate decisions in mouse ES cells. Biol Open 3:614–626. doi:10.1242/bio.20148409

Völkner M, Zschätzsch M, Rostovskaya M, Overall RW, Busskamp V, Anastassiadis K, Karl MO. 2016. Retinal Organoids from Pluripotent Stem Cells Efficiently Recapitulate Retinogenesis. Stem Cell Rep 6:525–538. doi:10.1016/j.stemcr.2016.03.001

Wegner M, Stolt CC. 2005. From stem cells to neurons and glia: a Soxist’ s view of neural development. Trends Neurosci 28:583–588. doi:10.1016/j.tins.2005.08.008

Winkler S, Loosli F, Henrich T, Wakamatsu Y, Wittbrodt J. 2000. The conditional medaka mutation eyeless uncouples patterning and morphogenesis of the eye. Development 127:1911–1919.

Xiao Y, Gao M, Gao L, Zhao Y, Hong Q, Li Z, Yao J, Cheng H, Zhou R. 2016. Directed Differentiation of Zebrafish Pluripotent Embryonic Cells to Functional Cardiomyocytes. Stem Cell Rep 7:370–382. doi:10.1016/j.stemcr.2016.07.020

Yi M, Hong N, Hong Y. 2010. Derivation and characterization of haploid embryonic stem cell cultures in medaka fish. Nat Protoc 5:1418–1430. doi:10.1038/nprot.2010.104

Zuber ME. 2003. Specification of the vertebrate eye by a network of eye field transcription factors. Development 130:5155–5167. doi:10.1242/dev.00723

